# O-glycans Expand Lubricin and Attenuate its Viscosity and Shear Thinning

**DOI:** 10.1101/2023.12.07.570567

**Authors:** Saber Boushehri, Hannes Holey, Matthias Brosz, Peter Gumbsch, Lars Pastewka, Camilo Aponte-Santamaría, Frauke Gräter

## Abstract

Lubricin, an intrinsically disordered glycoprotein, plays a pivotal role in facilitating smooth movement and ensuring the enduring functionality of synovial joints. The central domain of this protein serves as a source of this excellent lubrication, and is characterized by its highly glycosylated, negatively charged, and disordered structure. However, the influence of O-glycans on the viscosity of lubricin remains unclear. In this study, we employ molecular dynamics simulations in absence and presence of shear, along with continuum simulations, to elucidate the intricate interplay between O-glycans and lubricin and the impact of O-glycans on lubricin’s conformational properties and viscosity. We find the presence of O-glycans to induce a more extended conformation in fragments of the disordered region of lubricin. These O-glycans contribute to a reduction in solution viscosity but at the same time weaken shear thinning at high shear rates, compared to non-glycosylated systems with the same density. This effect is attributed to the steric and electrostatic repulsion between the fragments, which prevent their conglomeration and structuring. Our computational study yields a mechanistic mechanism underlying previous experimental observations of lubricin and paves the way to more rationally understanding its function in the synovial fluid.

## Introduction

Friction and adhesion involving viscous fluids or viscoelastic materials, such as those occurring in adhesives or lubricants, significantly impact various aspects of everyday life, from human interaction with the environment to the performance and reliability of machinery. Synovial joints, including the knee, hip, and shoulder, are essential for facilitating smooth movements of the human body. These joints exhibit remarkable lubricating properties, characterized by an extremely low coefficient of friction (ranging from 0.0005 to 0.04) and the ability to withstand pressures of approximately 200 atm (Figure 1A). Lubricin, a mucinous glycoprotein and intrinsically disordered protein (IDP), plays a crucial role in providing excellent lubrication by reducing friction within synovial joints. In the case of synovial joints, the viscoelastic response of lubricin at the molecular level gives rise to efficient tissue lubrication, enabling it to withstand high mechanical loads at the macroscopic scale. ^1–5^ However, the determining factors of lubricin’s molecular structure and dynamics underlying its lubricating function remain to be understood.

**Figure 1:**
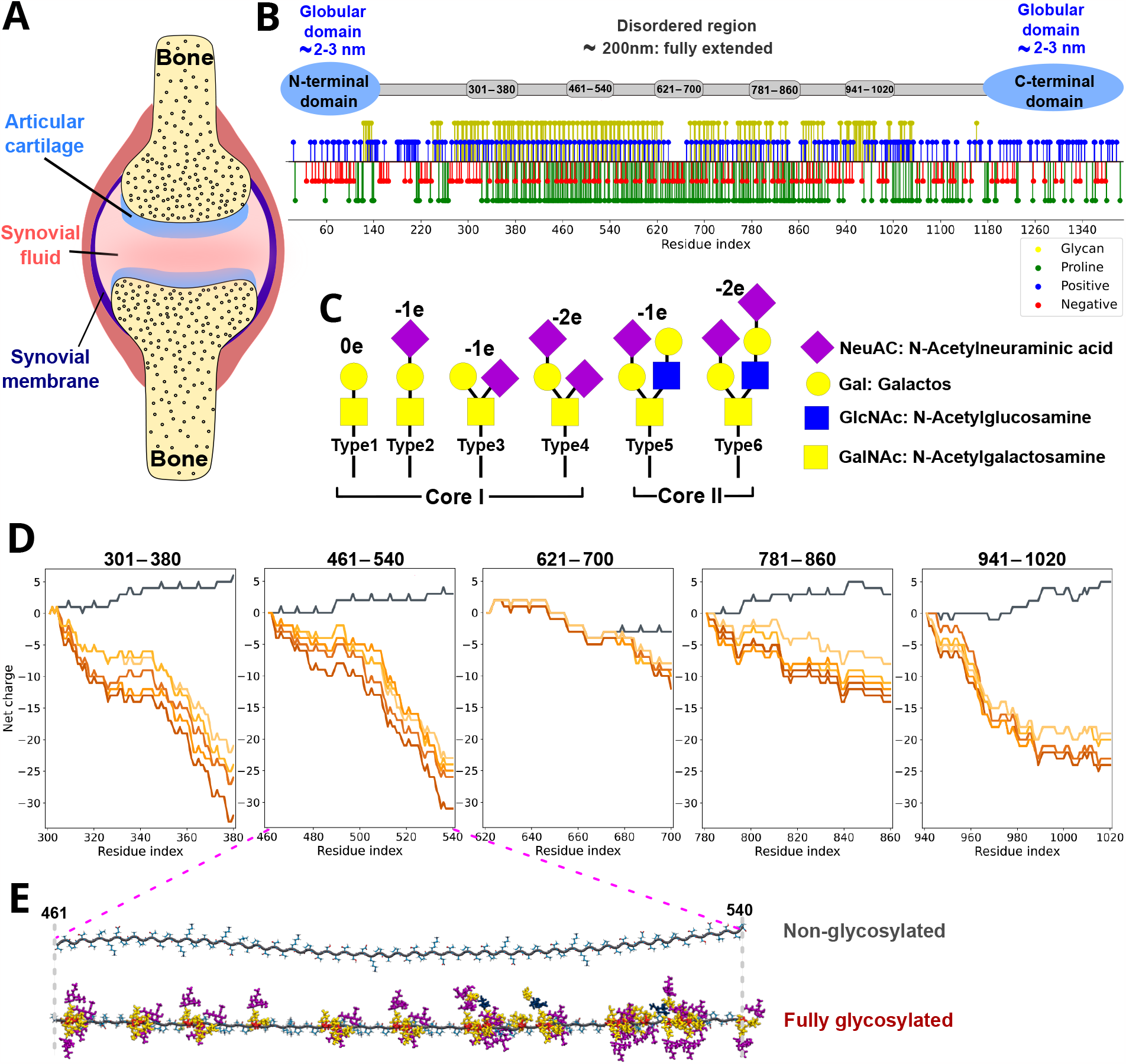
Sequence, charge, and glycosylation of lubricin. **A**. Schematic representation of the human synovial joint. Lubricin is found in the synovial fluid and acts as a lubricant to ensure the smooth movement of bones. **B**. Lubricin is composed of two globular terminal domains (blue) and a mucin-like central disordered region (grey). The two globular domains are non-glycosylated. The central region spans approximately 800 amino acids, is highly glycosylated by negatively charged O-glycans, and is rich in polar and proline residues. Glycan sites, charged residues, and proline residues along the sequence are indicated. **C**. Shown are Core I and Core II of six O-glycans of lubricin (out of eleven types) used in this study. The NeuAC sugar has a negative charge (purple diamond). The net charge of each type of O-glycan and the prevalence for lubricin are indicated. **D**. Five different 80-amino acid fragments spanning the disordered region of lubricin were considered. They were glycosylated to a different extent (six levels of glycosylation for each fragment). The cumulative net charge for each glycosylated fragment is shown. **E**. Examples of the initial fully extended structure for a non-glycosylated and a fully-glycosylated fragment (sequence 461–540) are shown.

Lubricin is encoded by the PRG4 (proteoglycan 4) gene. It is composed of approximately 1400 amino acids and has a length of approximately 200 nm ± 50 nm.^6^ Structurally, lubricin comprises two folded terminal domains, referred to as the N and C terminal domains, which are predominantly non-glycosylated, positively charged, and hydrophobic. These terminal domains enable lubricin to anchor to cartilage proteins, contributing to its ability to adsorb and withstand pressure.^7–9^ In contrast, the central domain of lubricin is overall intrinsically disordered.^10,11^ This region is known as the mucin-like domain and is highly glycosylated, negatively charged, and therefore highly hydrophilic. ^12,13^ Approximately 30-35% of the overall lubricin structure is composed of core I and core II O-linked glycans, namely sugar molecules attached to the hydroxyl group of serine (Ser) or threonine (Thr) residues. These O-glycans, along with sialic acid, contribute to negative charges in the protein^14–16^ (Figure 1 B, C). The mucin-like domain is very rich in prolines too. ^13^

The lubricating properties of lubricin are believed to be closely tied to its structural characteristics, with its high extent of glycosylation and disorder likely playing a vital role.^17,18^ However, the specific effects of glycosylation and disorder on the structure and viscosity of lubricin have yet to be fully elucidated. The synovial fluid shows shear thinning,^19–21^ just as many other industrial or biological systems, from molecular inks and Nafion to cellulose hydrogels, mucus, and blood.^22–25^ Shear thinning is one of the most common types of non-Newtonian behavior of polymers and characterized by a reduction in the viscosity of a fluid shear rate.^26^

Non-equilibrium Molecular Dynamics (MD) simulations have previously proven highly useful for studying the response of a polymer solution to shear at the molecular level. Shear rates can be induced across simulation systems by deforming the simulation box, using the so-called SLLOD algorithm,^27,28^ by dragging parallel walls relative to each other at a constant velocity,^25,29,30^ or also in combination with continuum models.^31,32^ Previous studies have successfully addressed the collapse propensities and intramolecular interactions of glycosylated disordered proteins by MD simulations at both atomistic^33,34^ and coarse-grained levels.^35^ However, their behavior under shear, to our knowledge, has not yet been explored.

In this study, we use a large set of equilibrium and shear-driven non-equilibrium MD simulations (Figure 2) and continuum calculations to explore the molecular determinants of lubricin’s function in reducing friction in synovial joints. Specifically, we aim to comprehensively understand how O-glycans impact lubricin-induced viscosity. We assess lubricin’s zero shear viscosity and its shear viscosities under varying shear rates.^25,36,37^ Glycosylation expands lubricin in equilibrium and reduces its intermolecular clustering. Glycosylation leads to overall lower viscosities when comparing the same mass densities, but these viscosities drop less pronouncedly under shear. Additionally, to demonstrate the broad utility of our molecular viscosity and shear thinning analysis in a multiscale context, we conduct representative continuum simulations using fluid models guided by our molecular dynamics data. ^38,39^ By shedding light on the structural and rheological properties of glycosylated lubricin, this study enhances our understanding of synovial joints.

**Figure 2:**
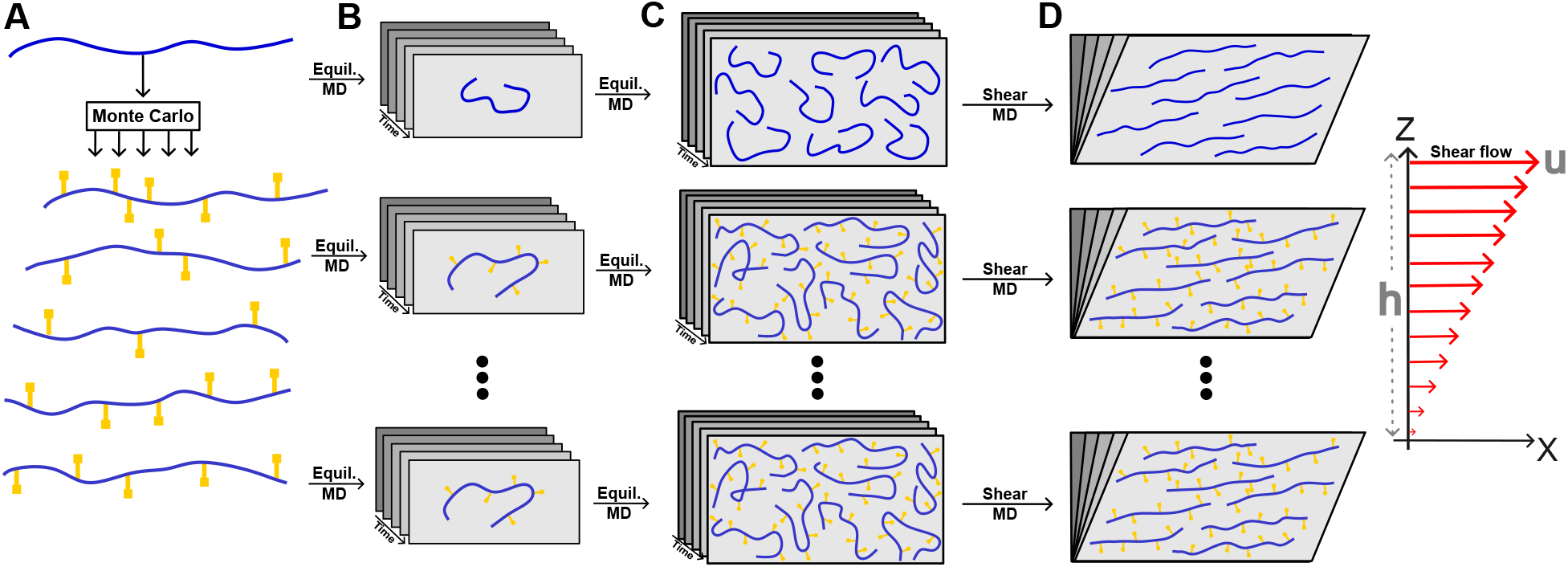
Molecular dynamics (MD) protocol to obtain viscosities of lubricin-derived fragments. **A**. By using Monte Carlo sampling, we generated five different glycosylation distributions for each of the five 80-amino acid long lubricin fragments. Here, the blue line indicates the non-glycosylated fragment and the yellow symbols depict the O-glycans. **B**. Equilibrium MD simulations were carried out for each fragment, both in its non-glycosylated and its five different glycosylated forms. **C**. Equilibrium MD simulations were performed for systems containing multiple fragments in order to obtain the viscosity in the absence of shear by using the Green-Kubo method. **D**. Shear-driven non-equilibrium MD simulations, deforming the simulation box, estimated viscosity under different shear rates.

## MATERIALS AND METHODS

### Glycosylation of lubricin fragments

In order to investigate the effect of O-glycans on the structure of lubricin, we selected five distinct segments, each spanning 80 amino acids of the central disordered part of lubricin (Figure 1B). These segments represent the protein’s physicochemical properties, including glycosylation, charged residues, and proline content. Among the 11 types of oligosaccharides that bind to lubricin,^15^ we chose six types (depicted in Figure 1C) based on their composition and charge (more information in Figure S1A). These six types of glycans were attached to the hydroxyl groups of glycosylated serine/threonine (Ser/Thr) side chains, as defined by Ali et al.,^13^ within each lubricin segment. The exact O-glycan binding at each position is unknown. For this reason, we added glycans, following five distinct oligosaccharide distributions based on a Monte Carlo sampling approach. In brief, we randomly added each type of glycan to the fragments based on the percentage (%) of total O-glycans (Figure S1B). By following this protocol, we generated one non-glycosylated and five different glycosylated versions of each lubricin fragment. The sugars that were attached at each glycosylation site, in the 25 different cases, are listed in Table S1-6, resulting in varying cumulative net-charge distributions (Figure 1D and Table S7).

We generated the initial structure of the non-glycosylated fragments by employing the Avogadro software package.^40^ The addition of O-glycans was carried out by using the glycam.org server. Fully elongated linear configurations were considered in all cases.

### Single-chain equilibrium MD simulations

Molecular dynamics (MD) simulations were performed using GROMACS (2020 version) software.^41^ The amber99sb-star-ildnp^42,43^ force field was employed for the protein, and the GLYCAM06^44^ force field for carbohydrates. Force field parameters from GLYCAM06 were converted to GROMACS format using the ACPYPE script.^45,46^

Initial structures were placed within a dodecahedron-shaped simulation box, solvated with TIP4P-D^47^ water molecules and 150 mM NaCl ions. Additional ions were added to neutralize the net charge of the fragments. The resulting systems had approximately 0.36 – 0.92 M atoms (see Table S8). The systems were subjected to energy minimization using a steepest descent algorithm until the maximum atomic force was below 1000 kJmol^−1^ nm^−1^. The systems were thermalized in the NVT ensemble at a temperature of 310 K by using the velocity rescaling thermostat^48^ during 1 ns (coupling time of 0.1 ps). Subsequently, the solvent was relaxed in the NPT ensemble at a pressure of 1 atm using the Parinello–Rahman barostat^49^ for 2 ns (coupling constant of 2.0 ps and reference compressibility of 4.5×10^−5^ bar^−1^). In both equilibration parts, a harmonic force (with an elastic constant of 1000 kJmol^−2^) was applied to restraint the position of the heavy atoms. Finally, production runs were executed under the NPT ensemble (same temperature and pressure as in equilibration steps) applying periodic boundary conditions and releasing the position restraints on the heavy atoms (Figure 2B). The velocity rescale thermostat and the Parrinello-Rahman algorithm were employed in these to maintain a constant temperature and a constant pressure during production runs too. Three replicas of 200 ns each were carried out for each of the 30 different fragments (see Table S8).

Electrostatic interactions were taken into account by using the Particle Mesh Ewald algorithm.^50^ Short-range interactions were modeled with a Lennard Jones potential truncated at a cutoff distance of 1.0 nm. Bonds involving hydrogen atoms of the glycosylated fragments were constrained by using LINCS.^51^ Accordingly, equations of motion were numerically integrated by using the Leap Frog algorithm at discrete time steps of 2 fs. Neighbors were treated with the Verlet Buffer with a tolerance of 0.005 kJmol^−1^ ps^−1^ and updating neighbors every 10 steps.

### Multi-chain equilibrium MD simulations

To investigate the influence of lubricin glycosylation on medium viscosity, five distinct lubricin systems were considered, each with the same molar and mass density (30 non-glycosylated, 73 non-glycosylated, 101 non-glycosylated, 30 medium glycosylated, and 30 highly glycosylated chains) (Figure 2C). The systems included 30 non-glycosylated (w.o. *ρ*=66.5 kg/m^3^) and 30 highly glycosylated (w.+ *ρ*=230.6 kg/m^3^) peptides, matched in molar concentration (N=30) but with different mass density. Subsequently, a system comprising 101 non-glycosylated peptides was introduced, maintaining a similar mass concentration to the highly glycosylated case (w.o. *ρ*=226.6 kg/m^3^). To approach a system more closely resembling reality, a system containing 30 randomly glycosylated peptides was selected (w. *ρ*=160. kg/m^3^). Furthermore, to investigate the impact of glycans, a system with 73 non-glycosylated peptides was considered to have the same mass density (w.o. *ρ*=158.6 kg/m^3^). Further simulation details can be found in Table S9.

The simulation box was filled with peptides adopting different conformations (taken from the single-chain MD simulations), TIP4P-D water molecules, 150 mM of NaCl, and extra neutralizing ions. A summary of the simulated systems and the resulting system sizes is presented in Table S9. Each system was equilibrated during 200 ns. Final conformations from this equilibration were considered for subsequent simulations under shear flows.

Additionally, two equilibrium MD simulations of 2000 ns each were conducted for each multichain system to estimate viscosity at zero shear (see below). The simulation parameters and equilibration protocol were identical to those used in the simulations of single chains (see Section Single-chain equilibrium MD simulations above).

### Zero Shear Viscosity

Due to computational constraints, non-equilibrium molecular dynamics (NEMD) simulations for very low shear rates were not feasible within the scope of our resources. To address this limitation, we calculated zero shear viscosity using equilibrium molecular dynamics simulations (EMD). In this study, the Green-Kubo method (GK)^52,53^ was utilized to estimate zero shear viscosity in the realm equilibrium MD simulation. The GK method allows for the determination of viscosity by integrating the autocorrelation function (ACF) of the pressure tensor components over time. Mathematically, viscosity (*η*) is determined through the following equation:^36,54^

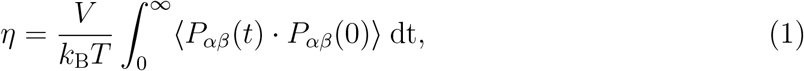

where ⟨*P*_*αβ*_(0)*P*_*αβ*_(*t*)⟩ is the correlation function of the *αβ* component of the pressure tensor (where *α* and *β* refer to x, y, and z coordinates and the brackets indicate ensemble average). Here, V, *k*_B_, T, and t denote the simulation box volume, Boltzmann constant, temperature, and time, respectively. Note that here we considered all six independent diagonal and off-diagonal components of the pressure tensor to calculate the ACF and thereby the zero shear viscosity: *P*_*αβ*_ = *P*_*xy*_, *P*_*xz*_, *P*_*yz*_, (*P*_*xx*_ − *P*_*yy*_)*/*2, (*P*_*xx*_ − *P*_*zz*_)*/*2, and (*P*_*yy*_ − *P*_*zz*_)*/*2.^55,56^

For estimating the zero shear viscosity, we focused on three specific systems: 30 non-glycosylated (w.o.), 30 medium glycosylated (w.), and 30 highly glycosylated (w.+) systems. The Green-Kubo method is acknowledged for its effectiveness primarily in fluids exhibiting relatively low viscosity, typically below 20 mPa·s.^57,58^ (Consequently, employing this approach to determine the zero shear viscosity for non-glycosylated systems with medium and high mass density (w.o. 73 and w.o. 101 chains) is challenging, due to their high expected viscosity.) Our approach consisted of conducting two prolonged simulations (2000 ns) for each system within the NPT ensemble. Subsequently, to ensure a comprehensive sampling of the viscosity of the systems, we extracted 25 conformations every 8 ns, from the last 200 ns of each of the two 2000 ns equilibrium simulations (50 extracted conformations in total). Each of these configurations underwent an initial equilibration phase of 2 ns in the NPT ensemble, followed by a subsequent simulation of 20-100 ns in the NVT ensemble to obtain zero-shear viscosity. Due to the high fluctuations of the pressure tensor to obtain a reliable estimate of the ACF, in these subsequent simulations, the pressure tensor components were written with a higher output frequency of 10 fs. Accordingly, the zero shear viscosity *η* was determined by averaging the results of the six distinct pressure components and over the estimates from the 50 distinct high-frequency short trajectories. The average was carried out after the viscosity converged in a time window (Figure 4A–C).

### MD simulations under shear flows

To further investigate the influence of shear flow on the viscosity of conglomerates containing lubricin fragments, we conducted shear-driven non-equilibrium MD simulations using the shear deformation method. The shear viscosity of the system can be determined by applying a Couette flow to the system.^59^ To generate a planar Couette flow, we considered the box deformation method and Lees-Edwards periodic boundary conditions^60,61^ (Figure 2D). In brief, the simulation box was deformed by moving the upper wall of the simulation box (i.e. located at *z* = *h*, with *h* the box size dimension along the *z*−axis) laterally at a constant speed *u* along the x-axis (Figure 2D). The *x*−component of the velocity of all particles contained in the simulation box was updated accordingly. This method provides a robust approach to generate planar Couette flow, with shearing occurring in the *x* direction while the velocity gradient goes along the *z* direction. In Couette flow, the velocity of the moving wall and the separation between the walls determine the shear rate 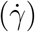:

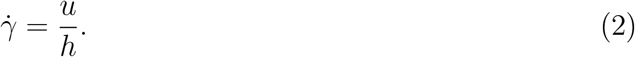

The shear viscosity (*η*) can be determined by evaluating the ratio between the component of the pressure tensor ⟨*P*_*xz*_⟩ and the shear rate of the fluid:

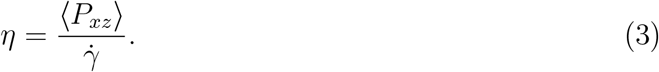

Here, we use the ensemble average of the component of the pressure tensor. The simulation box was deformed using a GROMACS-2023-dev version, with the Lees-Edwards boundary condition implemented. In these simulations, the pressure was not maintained constant and seven distinct deforming speeds were considered: *u* = 0.5, 1.0, 2.0, 4.0, 8.0, 10.0, and 15.0 nm/ns. The *h* value was equal to 15.9±0.1 nm in all systems. Accordingly, the resulting shear rates were: 0.03, 0.06, 0.13, 0.25, 0.51, 0.63, and 0.95 ns^−1^. Simulations of 50-600 ns in length were performed (Table S10), and the pressure tensor component was retrieved from these simulations.

### Viscosity of pure water

To validate our simulation protocol, we computed the viscosity of pure water. For the determination of zero shear viscosity, we used a box containing approximately 7100 TIP4P-D water molecules with dimensions of 6 × 6 × 6 nm^3^. Subsequently, for non-zero shear viscosities, a box with around 4300 TIP4P-D water molecules and dimensions of 8 × 4 × 4 nm^3^ was selected. The viscosity was computed at a temperature of 310 K, following the same simulation protocol as for the lubricin fragments, both at zero shear and under shear rates of 0.25, 0.50, 0.99, 1.99, and 2.48 ns^−1^. Simulation parameters and algorithms were identical to those used in the lubricin simulations, with the exception of the simulation time for non-zero shear rates, where simulations of 20-120 ns in length were performed.

### Simulation analysis

In addition to viscosity, the following observables were extracted from the molecular simulations. These quantities were computed as ensemble averages after discarding the first half of the simulations as equilibration time. Statistical uncertainties were reported as 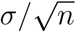, where *σ* corresponded to the standard deviation and *n* the number of independent samples.

### Radius of gyration

To assess the size of the different fragments, we utilize the radius of gyration. It was extracted from the equilibrium and non-equilibrium simulations by using the GROMACS tool.^41^

### Solvent accessible surface area

To monitor the aggregation between chains, we used the solvent-accessible surface area (SASA)^62^ as a quantitative measure. SASA measures the surface area of a protein or aggregation of proteins that is exposed to the surrounding solvent molecules. In our analysis, we calculated the value 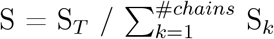, where S_*T*_ and S_*k*_ are equal to SASA of whole chains together and SASA of individual chain respectively. When S equals 1, it indicates complete dissociation of the chains, implying that they are not interacting with each other. Conversely, a low S value approaching 0 suggests a high degree of association among the chains, implying the formation of a condensed or aggregated state.

### Nematic order

The nematic correlation function (NCF) was computed to analyze the propensity of the chains to align with respect to each other. It delves into the orientation of individual polymer chains concerning each other. The NCF, typically denoted as Π_*ij*_(r), is calculated by considering pairs of amino acids, i and j, from two different protein backbone chains *I* and *J*, at a specific distance from each other within the system,^63^

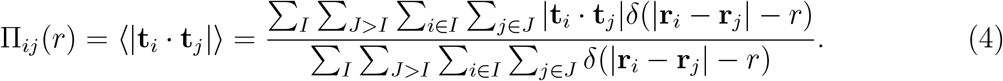

The value of Π_*ij*_(*r*) falls within the range of 0.5 to 1.0. A value of 0.5 indicates a complete absence of alignment, signifying that the orientation of fragments is entirely stochastic. Conversely, a value of 1.0 signifies perfect alignment, where the fragments exhibit a uniform orientation.

GROMACS analysis tools were utilized to extract the simulation observables. Further post-processing of the data was carried out by using in-house Python scripts. Trajectories were visualized and snapshots were rendered with VMD.^64^

### Continuum simulations

To highlight the transferability of our molecular characterization of viscosity and shear thinning in a multiscale approach, we performed exemplary continuum simulations with fluid models that are informed by our MD data. Therefore, we solve the one-dimensional steady Reynolds equation for an incompressible fluid

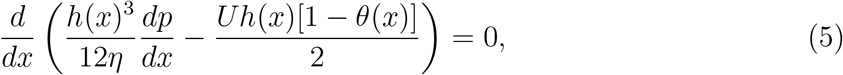

which is a simplification of the Navier-Stokes equation for laminar thin film flows including mass-conserving cavitation handled via the Jakobsson, Floberg, and Olsson (JFO)^65,66^ formalism. Equation 5 can be solved for the pressure distribution *p*(*x*) in a thin gap, given its height profile *h*(*x*) and a constant entrainment speed *U*. The cavity fraction *θ*(*x*) = 1−*ρ*(*x*)*/ρ*_0_ discriminates between full film (*θ* = 0) and cavitated regions (0 *< θ* ≤ 1), where *ρ*_0_ is the full film density. The Reynolds equation is accompanied by a complementarity constraint (*p* − *p*_cav_)*θ* = 0 to ensure that the pressure does not fall below a certain threshold *p*_cav_ (the vapor pressure) in the cavitated regions. We solve the so obtained linear complementarity problem with the Fischer-Burmeister-Newton-Schur (FBNS) algorithm,^67^ where the first and second term in Eq. 5 are discretized with a central difference and a first-order upwind scheme, respectively.

The Reynolds equation was originally derived for isoviscous fluids,^38^ but high pressures and shear rates in confined fluids often require the consideration of non-Newtonian effects such as piezoviscosity or shear thinning. Including these nonlinearities into the Reynolds equation is not always straightforward,^68^ but substituting the constant viscosity in Eq. 5 with an appropriate model 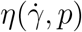 leads to good approximations in most cases. The solution to the nonlinear problem is then found via fixed-point iteration. Here, we employ the Carreau model for shear thinning^39^

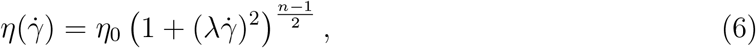

which describes the shear rate dependence of the stress as a power law with exponent *n <* 1 for 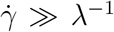 and approaches a Newtonian plateau at low shear rates. In our continuum simulations, we assume homogeneous shear thinning across the gap, i.e. the velocity profiles do not deviate from the Newtonian ones but the effective viscosity is reduced according to Eq. (6) with an average shear rate 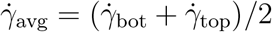, where 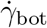 and 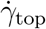 are the shear rate at the bottom and top wall, respectively. Note that this is a reasonable assumption only for weakly shear thinning fluids or where Couette flow dominates. For a more rigorous treatment of shear thinning application of the generalized Reynolds equation^69^ might be necessary.

## Results

### Glycosylation expands the conformation of lubricin fragments

Lubricin is a very large glycosylated protein, with a central disordered region of approximately 800 amino acids (Figure 1B). Simulating such a long, disordered protein is prohibitively expensive at the atomistic level of resolution. To elucidate the impact of O-glycans on the conformation of lubricin, we selected five 80-amino acid segments within the central disordered region (as depicted in Figure 1B). These segments vary in their number of glycosylation sites as well as contents of proline and charged residues. Subsequently, we generated five distinct glycan distributions for each of these segments by randomly attaching the six types of O-glycans according to their average proportions found in lubricin to all known O-glycosylation sites of the segment (Figure 1C, also see Methods).^13,15^ We repeated this procedure five times for each segment, which resulted, together with a non-glycosylated almost neutral state, in six different charge patterns per segment (Figure 1D,E). In a first step towards bulk lubricin solution systems for simulations under shear, we generated equilibrium structural ensembles considering single fragments, i.e. under infinitely-diluted conditions. We conducted multiple 200-ns MD simulation replicas for this purpose (Figure 2A,B). These fragments later served as starting points for assessing the viscosity of systems of solutions with high lubricin concentration (Figure 2C–D, next section).

To quantify the size of the fragments we computed the radius of gyration (R_*g*_). The glycans are bulky side chains that are expected to contribute to the overall radius of gyration of the fragments. However, to be able to assess the effect of glycosylation on the size of the fragments, we only considered the backbone atoms for the radius of gyration calculation. Starting from fully stretched conformations, within few tens of ns, the fragments partially collapsed, adopting conformations which ranged from approximately 2.3 to 3.9 nm (Figure S2). Figure 3A illustrates the correlation between the equilibrium R_*g*_ and the number of glycosylated residues per fragment. As expected, the more glycosylation sites are introduced, the more extended the conformation adopted by the fragments. The O-linked glycans attached to lubricin vary in composition and several of them carry a negative net charge (Figure 1C). This imposes an abundance of negative charge along the lubricin sequence (Figure 1D,E). Consequently, the size of the fragments increased by augmenting their net charge (Figure 3B), as previously observed for other IDPs.^70,71^ Note that the end-to-end distance or the solvent accessible surface area, which are other quantities indicative of the extension of the chains, consistently showed a similar trend as the radius of gyration (Figure S2). A linear fit of the data of the form *r*_*g*_ = *a*_*q*_|*q*| + *b*_*q*_ yielded a slope of *a*_*q*_ = 0.036 nm/e, which is about one fourth smaller than the value obtained in previous studies on IDP phosphorylation^70,71^ (i.e. 0.048 nm/e). The higher ionic strength of 150 mM used here (as opposed to 100 mM in^70,71^), by screening the electrostatic interactions further, may be responsible for the weaker response of the *r*_*g*_ to changes in charge observed here. In addition, the fit retrieved an intercept *b*_*q*_ = 2.72 nm, which is very close to the estimate for a random coil adapted to IDPs^72^ (R_*c*_ = *R*_0_ · *N* ^0.588^ = 2.61,nm, where *R*_0_ = 0.19 nm and *N* is the number of amino acids, specifically *N* = 80 in our case). In previous studies, the intercept was significantly smaller (1.7(1) nm^70,71^). The intercept is indicative of overall size of the chains under neutral conditions. In such case the bulky sugars, presumably acting as spacers, tend to promote more expanded conformations of the lubricin backbone fragments as compared as other IDP fragments with comparably smaller standard^70^ or phosphorylated^71^ amino acid side chains. Thus, O-glycans possess the capability to strongly expand the conformation of single-chain lubricin fragments. In the following, we investigated how lubricin influenced its medium’s viscosity.

**Figure 3:**
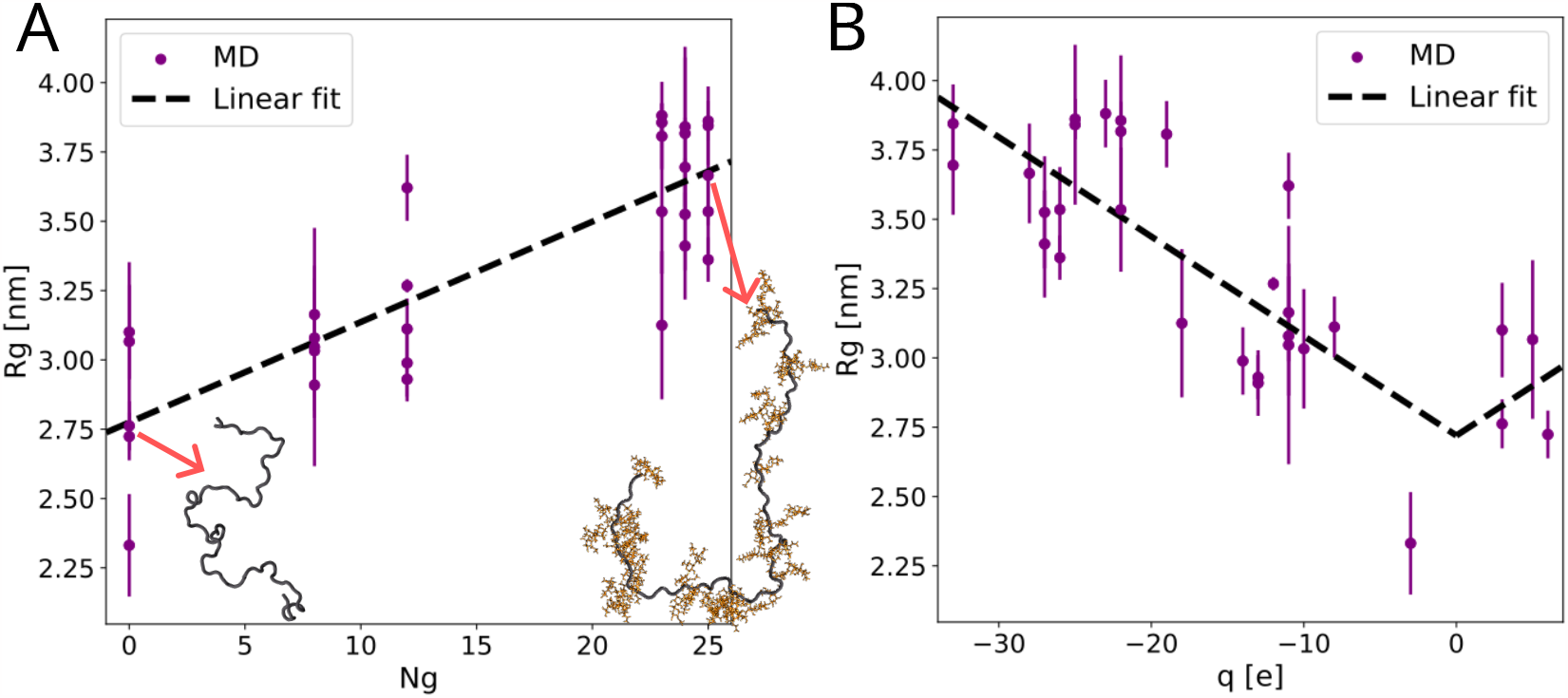
O-glycosyation increase extension and stiffness of lubricin’s single-chain fragments. **A–B.** Radius of gyration *r*_*g*_ as a function the number of glycosylated residues *N*_*g*_ (A) and the net charge of the fragment *q* (B). Symbols display the values recovered from the MD simulations (average ± s.e., *n* = 3). Dashed lines correspond to a linear regression of the data of the form *r*_*g*_ = *a*_*N*_ *N* + *b*_*N*_ (A), *r*_*g*_ = *a*_*q*_ |*q*| + *b*_*q*_ (B), with resulting fitting parameters *a*_*N*_ = 0.036 [1/e] and *b*_*N*_ = 2.77 [nm] (A) and *a*_*q*_ = 0.036 [nm/e] and *b*_*q*_ = 2.72 [nm] (B). Cartoons exemplify compact and extended conformations for a fragment without and with bound sugars, respectively (protein: black, sugars: orange).

### Viscosity

We next asked how O-glycans impact the rheological properties of lubricin. We calculated the viscosity of mixtures of fragments taken from the disordered region of lubricin in solution by conducting equilibrium and shear-driven non-equilibrium MD simulations (Figure 2C,D). Systems with different protein content (both in mass and molar density) as well as different levels of glycosylation were considered (see Methods and Table S9). Additionally, we computed the viscosity of pure water to validate our protocols. First, we employed the Green-Kubo method to calculate the zero shear viscosity from the autocorrelation of the pressure tensor (see Figure 4A–C and Methods section Zero Shear Viscosity) calculated from equilibrium MD simulations. Note that this method is primarily applicable to fluids exhibiting low viscosity, typically below 20 mPa·s.^57,58^ Consequently, it was not employed for the medium and high non-glycosylated systems, as the simulations with shear for these systems indicated a trend towards significantly larger zero-shear viscosities (see below). Secondly, the box deformation method and the Lees-Edwards periodic boundary conditions were used to obtain a Couette flow (Figure 4D,E and the method section MD simulations under shear flows). The resulting velocity profiles were linearly changing with the input shear rate (Figure 4E). Thus, this method enabled the utilization of equation 3 to estimate the viscosity of the system under different shear rates, based on the pressure along the shearing direction.

**Figure 4:**
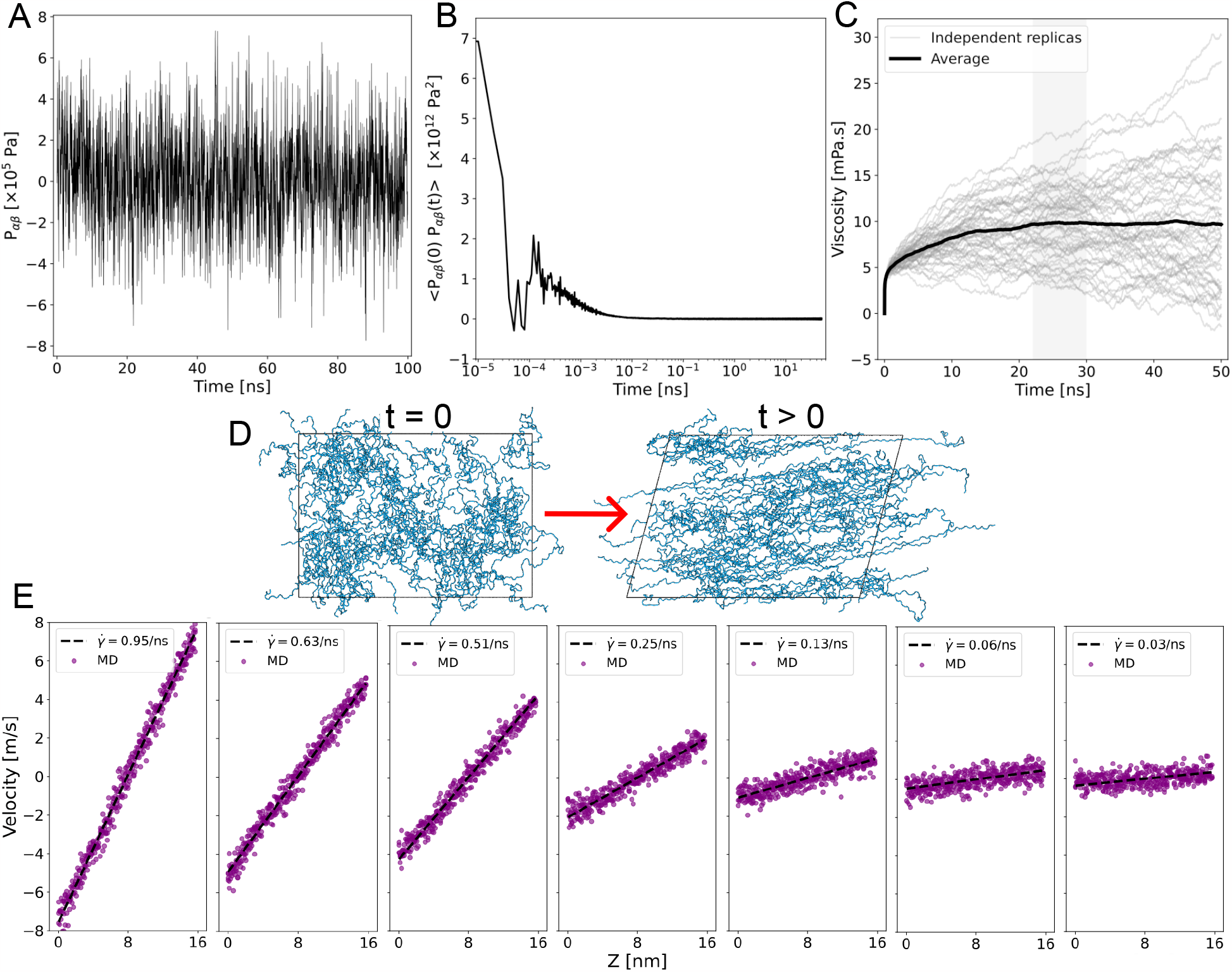
Viscosity calculation procedure at zero shear (A–C) and under shear flows (D–E). A. Components (*P*_*αβ*_) of the pressure tensor extracted from equilibrium MD simulations (shown here is the time-trace of one component for an exemplary system). **B**. The average of autocorrelation of all pressure components is computed for each independent system. **C**. The zero-shear viscosity is obtained from the average autocorrelation of all pressure components using equation 3 (see methods). Gray curves represent the independent viscosity of each replica (n=50). The black curve displays the average of all these curves. The viscosity was extracted from the plateau-highlighted region (average standard error). Figure S3 shows all three cases of zero shear viscosity. **D**. To estimate the shear viscosity, the simulation box was deformed at a shear rate 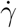. Initial (*t* = 0) and posterior (*t >* 0) snapshots are shown, highlighting the lubricin fragments in blue. **E**. Velocity profiles were obtained upon box deformation under different shear rates. The line corresponds to the linear fit. Output shear rates, i.e., *u/h*, are shown in the legend. Also, the expected values of shear rates are 0.94, 0.625, 0.50, 0.25, 0.125, 0.06, and 0.03 ns^−1^.

Figure 5A displays the resulting viscosities under different shear rates and at zero shear viscosity. As expected for a Newtonian fluid, the viscosity for pure water did not change in the presence of shear. Furthermore, the obtained values were close to the experimental value^73,74^ of 0.693 mPa.s (at the temperature of 310 K used throughout the simulations). Consequently, these results for water validate our simulation protocol and serve as a reference to examine the viscosity of systems with lubricin fragments.

**Figure 5:**
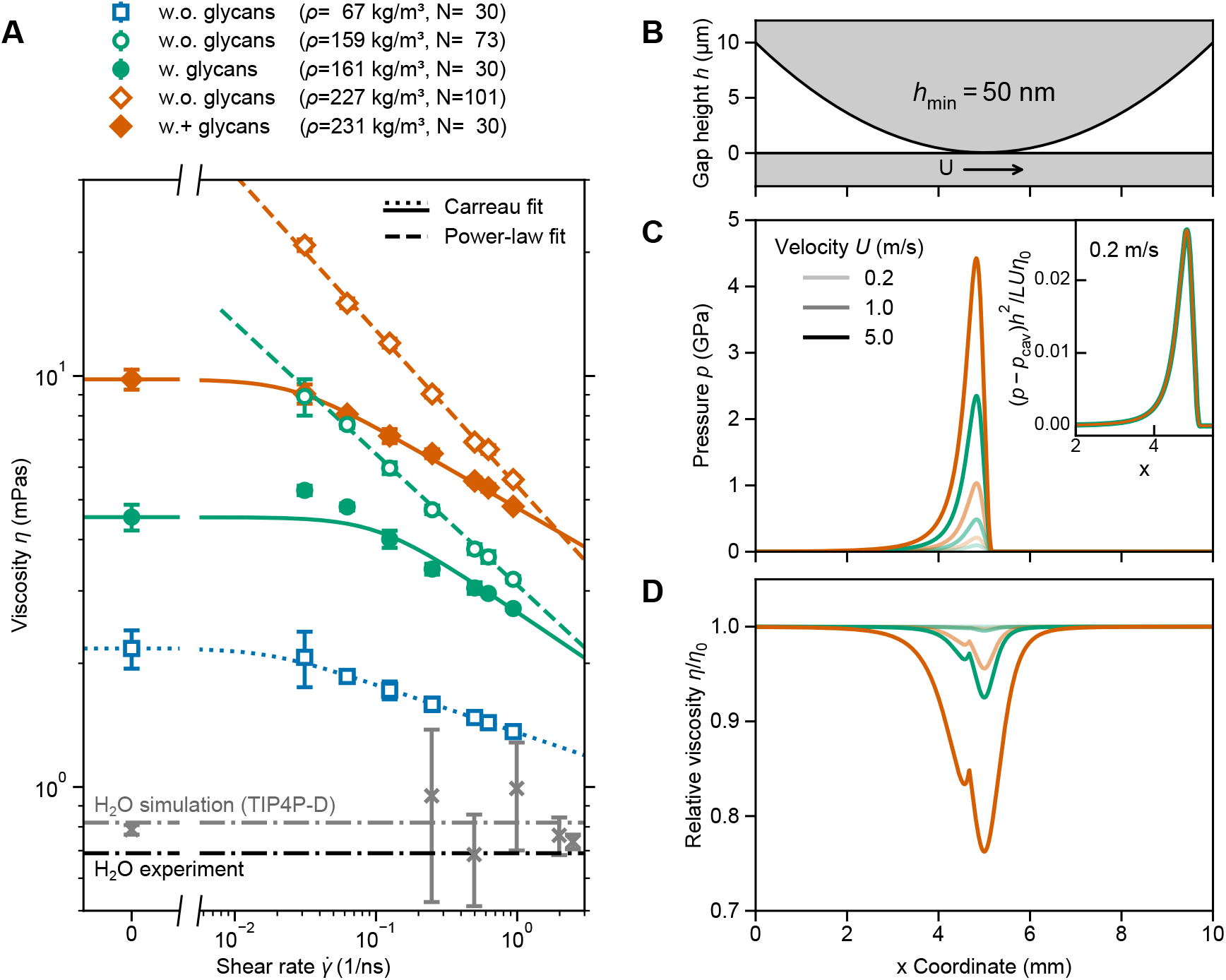
Shear thinning behavior of glycosylated lubricin fragments. **A**. Viscosity *η* as a function of the shear rate 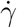 obtained from equilibrium simulations 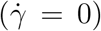 and shear-driven non-equilibrium MD simulations 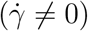. Five different systems were considered, either without glycans (“w.o”: open symbols and dashed or dotted lines) or with glycans(“w.”: closed symbols and solid lines). Systems with a medium (“w.”: green) and a high (“w.+”: orange) level of glycosylation were considered at the indicated mass (*ρ*) and molar densities (with *N* indicating the number of peptides in the system). A low-density system without glycans was also simulated (blue). For systems where zero-shear Newtonian viscosities are available, fits to the Carreau model (Eq. 6) are shown (solid or dotted lines). Dashed lines are fitted to a simple power law expression 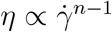 without Newtonian plateau. The viscosity of water, both experimental and obtained here, is also displayed as reference (grey and black, respectively). **B**. Sample geometry for Reynolds calculations with a flat wall sliding at velocity *U* against a parabolic height profile *h*(*x*) = *h*_min_ + 4(*h*_max_ − *h*_min_)(*x/L*_*x*_ −1*/*2)^2^ with *h*_min_ = 50 nm, *h*_max_ = 10 *μ*m, and *L*_*x*_ = 10 mm. **C**. Pressure profiles for lubricants with a medium and high level of glycosylation and three different sliding velocities. At low speed, normalized pressure profiles (by a reference pressure *η*_0_*LU/h*^2^) fall onto the same curve, indicating that the flow is still in the Newtonian regime, as shown in the inset. **D**. Effect of shear thinning in the Reynolds calculations shown by the local relative viscosity 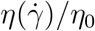 along the sliding direction.

The systems containing lubricin fragments featured higher viscosities than the system composed of only water. Unlike pure water, the viscosity of these systems changes with each shear rate, and with increasing shear rate, the calculated viscosity decreases slightly, indicating that these systems transition to a non-Newtonian regime exhibiting shear thinning behavior (Figure 5A). The change in viscosity with shear varied drastically among the different systems. Reduction in the protein mass density decreased the viscosity, approaching the value observed for pure water (compare curves for different densities, i.e. different colors). A further reduction in sensitivity to shear (i.e. the slope in the viscosity-shear curve) was observed for the glycosylated systems compared to their non-glycosylated counterparts (compare non- and glycosylated systems, i.e. dashed with solid lines). Within the same mass density, the glycosylated systems (solid lines) exhibited lower viscosity compared to non-glycosylated (dashed lines) counterparts.

A Carreau model (equation 6) explains well the viscosity-shear relationship obtained for the systems for which the zero-viscosity was computed (Figure 5A and Table 1). However, such an analytical model could not be applied to the systems lacking zero-shear viscosity, namely the medium and high-density non-glycosylated ones (Figure 5A). A simple power law expression, without the Newtonian plateau, was fitted in these cases instead, and it also explained the simulation data well (Figure 5A and Table 1). The Carreau fit retrieved zero-shear viscosities that were largely dependent on the protein density. Note that although the medium- and high-density non-glycosylated systems lacked a direct estimate of the zero shear viscosity, extrapolation of their viscosity towards low shear rates suggests the zero-shear viscosity to take even larger values for these two systems than the glycosylated ones (Figure 5A). The shear rate at which the system transitioned from a Newtonian to a non-Newtonian regime is related to 1*/λ* (see equation 6). This value was of the order of 10^−2^–10^−1^ ns^−1^. Both models predicted an exponent *n* which measures the extent of decrease in viscosity with shear rate, where small values of *n* indicate strong shear thinning. These values are comparable across systems, although they are consistently higher for the glycosylated systems (compare *n* for systems with and without glycans in Table 1).

**Table 1:** Fitting analytical models to the viscosity-shear curves obtained from MD simulations. Five different systems were considered, either without glycans (“w.o”) or with glycans(“w.”), at the indicated mass (*ρ*) and molar densities (with *N* indicating the number of peptides in the system). For systems where zero-shear Newtonian viscosities were available, the Carreau model (Eq. 6) was applied, yielding three fit parameters: zero-shear viscosity *η*_0_, the exponent *n*, and the scaling factor *λ*. For systems lacking zero-shear viscosities (–) a simple power law expression 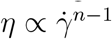 retrieved the parameter *n*.

| System | $\rho$<br>(kg/m<sup>3</sup>) | $N$ | $\eta_0$<br>(mPas) | $n$<br>(–) | $\lambda$<br>(ns) |
| --- | --- | --- | --- | --- | --- |
| w.o. glycans | 66.5 | 30 | 2.173 | 0.884 | 56.7 |
| w.o. glycans | 158.6 | 73 | – | 0.681 | – |
| w. glycans | 160.5 | 30 | 4.535 | 0.766 | 9.7 |
| w.o. glycans | 226.6 | 101 | – | 0.622 | – |
| w.+ glycans | 230.6 | 30 | 9.813 | 0.802 | 38.1 |

In summary, our data demonstrates that, in the high shear-regime, a pronounced shear thinning behavior of lubricin mixtures and its strong modulation exerted by the presence of O-glycans.

### Effect of shear thinning in macroscopic simulations

To show possible applications of the shear thinning models to macroscopic systems, we performed continuum simulations that highlight the effect of lubricin’s viscosity on pressure profiles generated in a lubricated sliding contact. We focused on a simplified setup that consists of a parabolic gap height profile sliding at speed *U* against a flat wall (although we have chosen the frame of reference to be located on the profiled surface for simplicity, see Fig. 5B). Furthermore, we assumed that the profile extends infinitely into the direction perpendicular to the sliding velocity and the gap height.

The solution to the Reynolds equation 5 provides pressure profiles that are generated in front of the geometrical constriction as shown in Fig. 5C). For a given geometry, the pressure excursions depend on viscosity and sliding speed only. Here, we compare the medium (green) and highly (orange) glycosylated configurations for three different sliding speeds *U* ∈ [0.2, 1.0, 5.0] m*/*s as indicated by the level of line opacity. It is not surprising that increasing either sliding velocity or viscosity (by increasing the level of glycosylation) leads to higher pressures. Behind the constriction, the fluid film cavitates and the pressure equals the cavitation pressure. The load bearing capacity, i.e. the force per unit length theoretically required to maintain the pre-defined gap height distribution (without considering elastic deformation of the walls) reaches 2.6 kN*/*mm for the highest pressure excursion (highly glycosylated and *U* = 5 m*/*s).

Looking at the pressure profiles alone does not elucidate the effect of shear thinning. In the Newtonian, i.e. linear response, regime, normalized pressure profiles fall onto the same curve, which was only approximately true for the lowest sliding velocity as shown in the inset of Fig. 5B. In contrast, the highest sliding velocity led to non-Newtonian flows as illustrated in the viscosity profiles along the contact line in Fig. 5D. We observed viscosity reduction down to less than 95% and 80% of the Newtonian viscosity at the point where the magnitude of the pressure gradient is largest for the medium and highly glycosylated system, respectively. Direct comparison of shear thinning for glycosylated and non-glycosylated peptides with our Reynolds calculations is difficult, because our MD data does not resolve the Newtonian plateau for the latter. However, we expect a much higher shear thinning for systems without glycans, as reflected by the exponents from the fits to the Carreau and the power-law models, see Tab. 1. In the shear thinning regime, a tenfold increase of the shear rate reduces the viscosity by 37 % in the system with high level of glycosylation (*n* = 0.80), compared to 58 % in the system without glycans (*n* = 0.62).

### Rheology of Lubricin under shear stress

We next investigated the molecular causes that gave rise to the observed shear thinning behavior of the systems containing mixtures of lubricin fragments. We analyzed distinct structural attributes within these mixtures related to the elongation, aggregation, and alignment response of the lubricin fragments under the influence of external shear stress.

We first analyzed the elongation of individual chains within the mixture by computing their radius of gyration (R_*g*_). Figure 6A displays the impact of shear stress on the R_*g*_. In all cases, an increase in shear rate induced an increase in the R_*g*_, indicating that the chains adopt more elongated conformations under shear stress. The reference value at zero-shear displayed a strong dependency on mass density and glycosylation: as the mass density increased, glycosylated un-sheared chains were found to be more elongated, while the non-glycosylated un-sheared ones were more compact (compare zero-shear values for different mass densities and glycosylation in Figure 6A). The elongation response under shear was also dependent on these two factors. Sheared glycosylated chains were found to be more elongated as the mass density increased, an effect that could be attributed to the electrostatic repulsion between the negatively-charged sugars. Surprisingly, non-glycosylated chains also displayed a similar trend, despite lacking such repulsion (see the increasing offset with density in the elongation curves in Figure 6A). However, overall, the elongation of glycosylated chains significantly surpassed that of non-glycosylated counterparts for each mass density.

**Figure 6:**
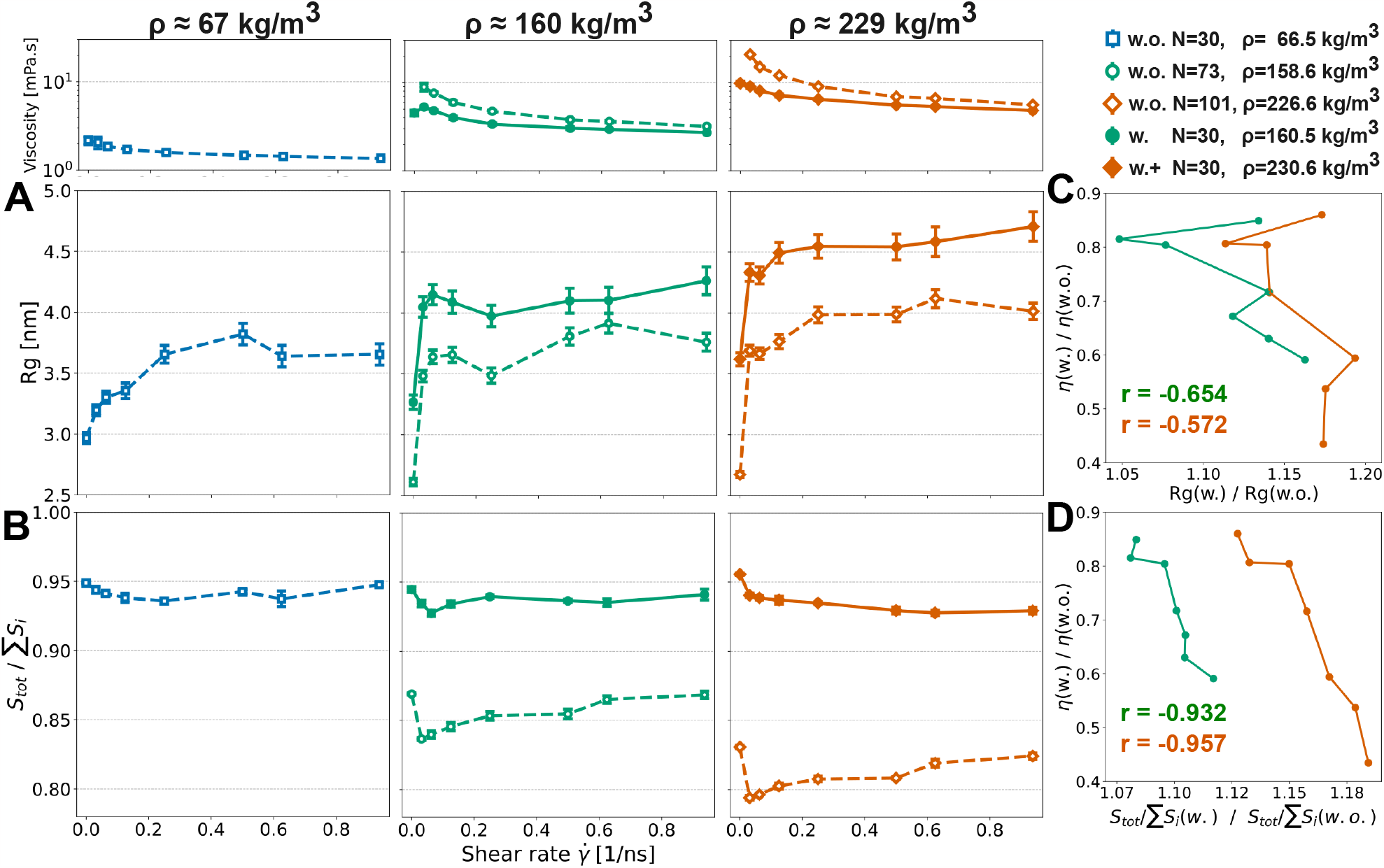
Shear-dependent rheological properties of lubricin fragments derived from MD simulations. **A-B**. Variation of the radius of gyration (R_*g*_) (A) and the ratio between the SASA of whole proteins and the cumulative SASA of individual proteins (surface exposure ratio, see methods section Solvent accessible surface area) (B) as a function of the shear rate 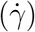. Low values of S_*tot*_*/Σ* S_*i*_ indicate a high degree of protein aggregation whereas a value of one denotes zero aggregation. Symbols represent data obtained from Equilibrium Molecular Dynamics simulations at zero shear rate and shear-driven non-equilibrium MD simulations at non-zero shear rates (average standard error, *n* = 4). Dashed lines correspond to non-glycosylated systems (‘‘w.o.”), while solid lines represent glycosylated ones (medium glycosylated: ‘‘w.” and highly glycosylated: ‘‘w.+”). Color indicates mass densities. The viscosity shear-response (of Figure 5A) is shown at the top of A for comparison. **C-D**. The ratio of viscosity *η*(*w*.)*/η*(*w*.*o*.) is presented as a function of the ratio *X*(*w*.)*/X*(*w*.*o*.), with *X* =R_*g*_ (C) and *X* = S_*tot*_*/Σ* S_*i*_ (D), in both the medium (green) and high (orange) mass density regimes. Correlation coefficients (r) for each data set are indicated.

We next investigated the aggregation tendency of the chains by computing the solvent-accessible surface area (SASA) of the whole protein conglomerate divided by the sum of the SASA of each individual chain. Accordingly, this ratio takes a maximum value of one when the chains are completely separated, while it diminishes when the chains aggregate (see methods section Solvent accessible surface area). Figure 6B shows the impact of shear flow on the interaction behavior of the lubricin mixtures. In all instances, imposing shear stress results increase intermolecular interactions between fragments relative to their state under zero shear rates (compare zero versus non-zero values in Figure 6B). Nevertheless, glycosylated and non-glycosylated mixtures displayed a distinct response to shear. Inter-chain interactions of glycosylated mixtures was overall low and practically insensitive to shear and to mass density (see high SASA ratio and practically constant values for different shear and density conditions). On the contrary, non-glycosylated chains displayed a strong level of interactions (low SASA ratio) directly proportional to mass density but inversely related to shear (see a monotonic increase in the SASA ratio with shear with an augmenting offset as density increased for the non-glycosylated chains).

The third structural attribute that we investigated was the alignment of the lubricin chains, assessed by computing the Nematic Correlation Function (NCF). This function monitors the correlated orientation of all chains weighted by their radial distribution function (see methods section Nematic order). Figure 7A illustrates that across all scenarios, higher shear rates lead to a stronger alignment of chains. Higher shear rates enhance chain alignment for both nearest neighbor chains (Figure 7B) and distant chains (Figure 7C), as also visible from examplary snapshots shown in Figure 4B. Note that due to the bulky glycan side chains, the distance to the first aligned neighbor was found to be higher for the glycosylated systems, indicating more spacing between these chains (Figure 7A). This is another indication of the low level of interactions for this case (Figure 6B). Within non-glycosylated systems, increasing the mass density results in a greater degree of alignment (Figure 7A), both locally for the first-neighbor chains (Figure 7B) and for distant chains (Figure 7C). Conversely, in glycosylated systems, alignment is less sensitive to changes in density (Figure 7A). Glycosylated chains align in a less correlated fashion with their nearest neighbors and pretty much equally with the distant ones, compared to the non-glycosylated ones (compare glycosylated versus non-glycosylated cases in Figures 7B,C).

**Figure 7:**
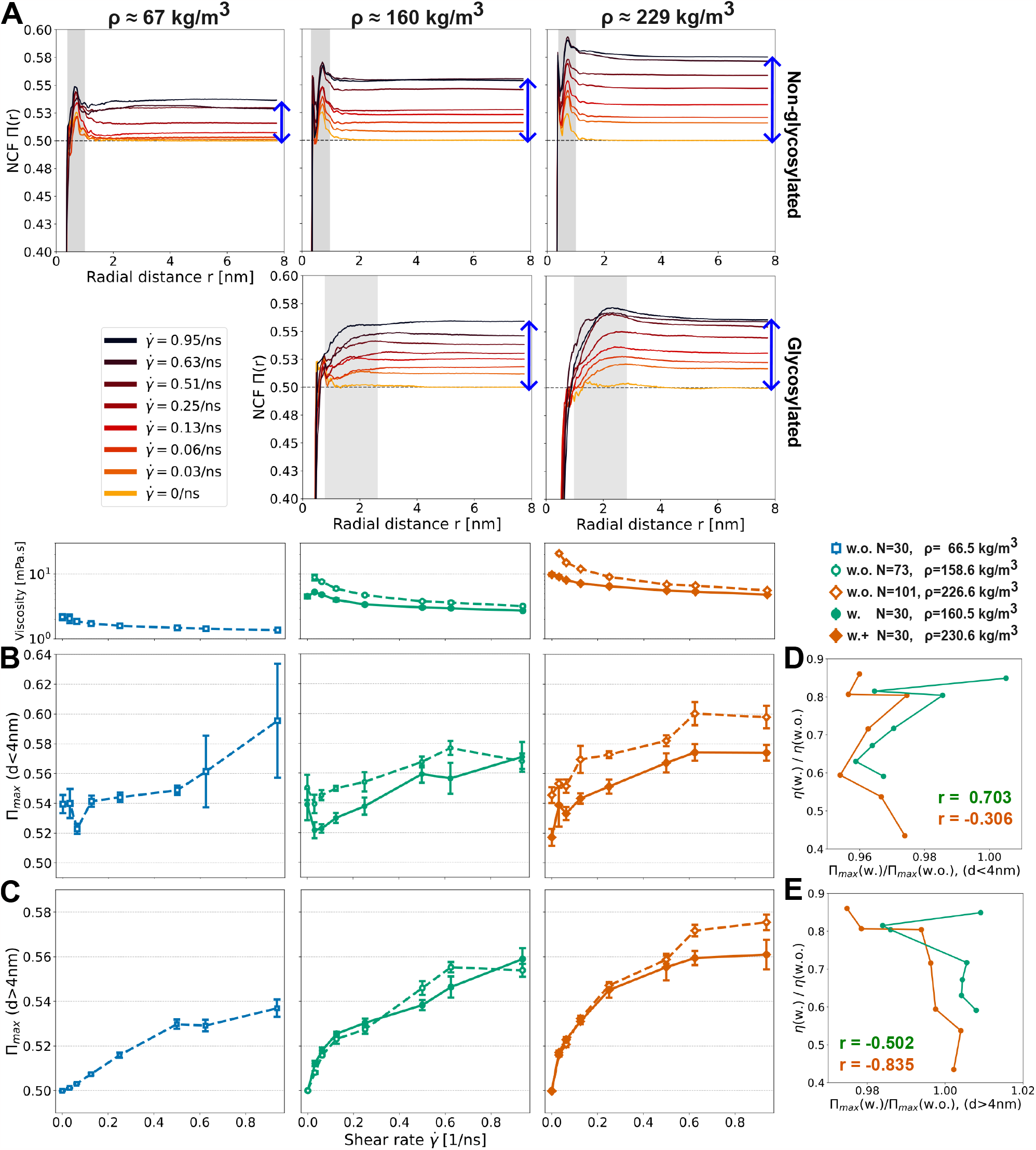
Shear-dependent alignment of lubricin fragments derived from MD simulations. **A**. The NCF for bulk systems of non-glycosylated (top row) and glycosylated (bottom row) systems with different mass concentrations (columns) as a function of the radial distance. Color indicates the different shear rates. An NCF value of 0.5 indicates random chain orientation while 1.0 indicates full alignment. The gray area corresponds to nearest neighbors region for which high alignment was observed. **B-C**. The NCF for short range (<4nm) (B) and for long range (>4nm) (C) inter-chain separations as a function of the shear rate 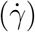. Symbols depict data obtained from the simulations (average ± standard error, *n* = 4). Dashed lines correspond to non-glycosylated systems (‘‘w.o.”) while solid lines represent glycosylated ‘‘w.”systems. Colors indicate mass density. The viscosity shear-response (of Figure 5A) is shown at the top of B for comparison. **D-E**. The ratio *η*(*w*.)*/η*(*w*.*o*.) is presented as a function of the ratio *X*(*w*.)*/X*(*w*.*o*.), with *X* = *NCF*_max_, i.e. the maximum of the NCF, at short-range (C) and the long-range (D) separations, in both the medium (green) and high (orange) mass density regimes. Correlation coefficients (r) for each data set are indicated.

To connect these structural characteristics with viscosity, we computed the ratio of viscosity between glycosylated and non-glycosylated systems and compared it with the corresponding ratio of the radius of gyration, solvent-accessible surface area ratio (Figure 6C,D), and nematic order parameter (Figure 7D,E). This analysis suggests that the divergent shear thinning trend observed between glycosylated and non-glycosylated systems is more closely correlated with inter-chain interactions, that is, clustering of chains, than with their relative ordering (see higher correlation in Figure 6D than in Figures 6C and 7D,E).

In summary, the structural response of conglomerates to shear is tightly controlled by both the protein density and the extent of glycosylation.

## Discussion

By using atomistic MD simulations under shear, here we investigated the role of O-glycans on the structural and rheological properties of lubricin fragments. A central finding of our study is that the viscosity and shear thinning behavior of mixtures containing short fragments derived from lubricin are tuned by the level of O-glycosylation of these fragments.

The conformational dynamics of IDPs is tightly regulated by post-translational modifications, such as phosphorylation, ^70,71,75^ methylation^76^ and glycosylation.^33,77^ Lubricin contains a large disordered region which is highly glycosylated (Figure 1B). We here show glycosylation strongly influences the structural properties of single 80 AA fragments distributed across this disordered region. A notorious difference of glycosylation with other post-translational modifications is the size of the attached moiety. While in phosphorylation or methylation, small moieties are added to side chains, in glycosylation, large and bulky glycans are attached to the protein. This is the case for lubricin which displays a complex and dense glycosylation pattern^33,77^ (Figure 1C). Accordingly, glycosylation promoted more expanded conformations for lubricin fragments than those observed for other IDPs of comparable sequence length^70,71^ (Figure 3). Nevertheless, the size of the fragments linearly increases with their net charge following a similar V-pattern.^70,71^ This emphasizes the important role electrostatics have governing the conformation of IDPs and the ability of post-translational modifications to tune such conformations. In the case of lubricin, the sensitivity of the conformational ensemble to glycosylation appears to be a crucial determinant of its viscosity response to shear (as discussed below).

To assess the viscosity response to shear, we employed equilibrium and shear-driven non-equilibrium MD simulations (Figures 2 and 4). As an alternative and more-affordable computational approach to examine the effect of lubricin and, more specifically, its glycosylation on the medium viscosity, we avoided the consideration of the full ∼800 aminoacid long disordered region, but instead a mixture of short glycosylated fragments extracted from it. This constitutes a first attempt to resemble the lubricin content in a small volumetric unit. To validate our protocol, we calculated the viscosity of pure water. Importantly, we observed its Newtonian behavior, namely, we obtained a viscosity that is independent of the applied shear (Figure 5A). Moreover, we obtained values which are consistent with the experimental viscosity of water^78^ and with the viscosity estimate for the TIP4P-D water model,^47^ the water model suited for IDPs which has been used in this study. To obtain the viscosity under shear, we performed shear-driven non-equilibrium MD simulations in which the box was deformed according to the Lees-Edwards periodic boundary conditions, as previously suggested by.^61^ We made use of a developmental implementation of such methodology in the GROMACS package^41^ (GROMACS-2023-dev version). We confirmed that, in fact, this methodology can retrieve sustained planar Couette flows (Figure 4D–E) and can be used to reliably estimate the viscosity of the medium. Consequently, this approach becomes attractive for studying the shear response of other biomolecular systems with the widely-used GROMACS package. Note, however, that large deformation velocities had to be applied. We opted for the Lees-Edwards boundary conditions as opposed to two walls moving relative to each other to induce the shear,^60,61^ as the latter setup requires appropriate description of the wall-solvent and wall-solute interactions. Such walls are absent in the biological or experimental setting and can induce artefacts. To close the gap between experimental observations, typically at low or zero shear,^79^ and our shear-driven non-equilibrium MD simulation results, we also computed the viscosity at zero shear, from the autocorrelation of the pressure tensor components, by using the Green-Kubo method^36,80^ (Figure 4A–C). However, this choice excluded systems exhibiting large viscosities, i.e. *>*20–30 mPa.s.^57,58^ Medium- and highly-dense non-glycosylated lubricin systems precisely fell into this category but they still served as reference of the overall shear-response when glycans were not present. In addition, with the help of the Carreau model (equation 6) we could predict the viscosity for the whole range of shear, and in its absence too, for the glycosylated systems.

Parameterization of semi-empirical viscosity models such as the Carreau model can be leveraged in multiscale simulations of lubrication. By doing so, molecular mechanisms to accommodate externally applied shear can be effectively transferred to length and time scales initially inaccessible to MD simulations. The simulation setup in Fig. 5B has been chosen without having a specific application in mind and should be primarily seen as a proof of concept. However, it is easy to imagine that the gap height profile could be replaced by the actual topography of an AFM tip or macroscopic synovial joint replicates. Recent macroscopic simulations of synovial joint lubrication, such as those considering pressure-induced rehydration by cartilage surfaces,^81^ might benefit from MD-informed constitutive relations. Similarly, other effects such as pressure-dependent viscosity, compressibility, or localized shear might be subsequently introduced into hydrodynamic descriptions. ^82^

Lubricin is believed to act as a lubricant in the synovial fluid.^2–6^ Ludwig et al. reported viscosities in the range of a few mPa.s. ^79^ Encouragingly, our estimates for glycosylated systems at zero shear are in the same order of magnitude (Figure 5A). Note, however, that for rendering our simulations computationally feasible, we had to use a much higher density (from 60 to 230 kg/m^3^, i.e. at least 500 fold higher than the experimental protein density^79^) and had to consider shorter protein chains. The experimental viscosity seems to have saturated at even lower densities, as moving from 0.0045 to 0.450 *kg/m*^3^, i.e. 100-fold, leads to only small change of ∼ 1 mPa.s. Our results are consistent with this behavior as a 500-fold higher protein concentration, used in our simulations, predicted viscosities in the same order of magnitude.

Our simulations together with the Carreau model predict shear-thinning to occur at high shear rates (*>* 10^−2^*/*ns i.e. *>* 10^7^*/*s) and a Newtonian behavior otherwise, for both studied glycosylated systems and the non-glycosylated system at low density (Figure 5 and Table 1). In agreement, at experimentally accessible rate regimes(∼10^0^–10^3^/s), pure glycosylated lubricin exhibits a Newtonian behavior.^79^ Shear rates are believed to vary widely in the synovial joint, reaching *>* 10^6^*/*s,^83^ and may even exceed this range on the microscopic scale e.g. due to surface roughness. The role of shear thinning at these larger shear rates as observed in our simulations is currently unknown. Larger simulation systems at coarser resolution will be required to more directly address the rheology in the physiological setting. Such systems can accommodate longer lubricin chains and other synovial components (such as hyaluronic acids) known to impact fluid viscosity,^79^ and incorporate boundary effects such as surface attachment.^21^

It is remarkable that despite its simplicity, our model seems to contain the necessary molecular features that give rise to observed lubricin medium’s viscosity. We suspect the key feature is the inclusion of glycosylated stretches from the disordered region (Figure 1). In fact, O-linked glycosylation was found to critically modulate the shear thinning response and the value to which the viscosity approached at zero shear (Figure 5A). The glycosylation status exerts a significant influence on structural parameters that dictate the fragment’s elongation (Figure 6A), interactions (Figure 6B) and alignment (Figure 7). Remarkably, the presence of glycans prevents formation of conglomerates, presumably due to the steric and electrostatic repulsion between fragments. This leads to greater inter-fragment spacing, where water can accommodate, and to reduced intermolecular interactions. In contrast, non-glycosylated systems, after experiencing extension due to shear stress, establish conglomerates with reduced spacing available for water and pronounced intermolecular interactions. Note however that such systems eventually disperse in response to external shear stress. The strong drop of viscosity in unglycosylated lubricin simulations is dampened upon glycosylation as they largely lack interchain interactions and thus are less affected in their structural ensemble by shear (Figure 6D). Interestingly, neither the elongation (Figure 6C) of the chains nor their alignment (Figure 7D,E) directly relate with the dampening. Overall, the lowering in viscosity by shear is suggested to be related to the amount of space between chains available to the solvent, a quantity that is promoted by the shear-dependent elongation and alignment of glycosylated chains. It is likely that other highly glycosylated systems such as mucin or aggrecan show a similar behavior when sheared.^84,85^

Taking these observations together, multi-site O-glycosylation affects the viscosity two-fold: First, it renders the initial (zero-shear) viscosity relatively low due to reduced inter-chain interactions, compared to a non-glycosylated lubricin system at the same mass density. Secondly, it lets the viscosity fall of less dramatic at higher shear rates, maintaining values far above pure water. Lubricin solutions thus are ‘robustly viscous’, being able to nearly maintain their moderate viscosity over many orders of magnitude of shear rates, very different to their unglycosylated counterparts, which at drastic shear loads can not uphold a significant resistance.

## Conclusion

Our study aimed to investigate the impact of glycosylation on the structure and viscosity of lubricin, a vital protein involved in facilitating lubrication within synovial joints. Through extensive molecular dynamics simulations and continuum simulations we gained valuable insights into the role of O-glycans in influencing lubricin’s properties.

Our findings reveal that glycosylated disordered lubricin solutions, on one hand, exhibit lower viscosities at zero shear, thus lower friction, when compared to unglycosylated proteins at the same mass density. On the other hand, glycosylation attenuates the pronounced loss of viscosities with shear rates exhibited by their non-glycosylated counterparts. According to our results, the molecular basis of this effect is the presence of O-glycans, which promote the dispersion of lubricins thereby preventing their clustering. Our continuum model showcases how quantitative viscosity relations from nanometer-scale molecular simulations can be harnessed to draw conclusions on much larger length scales and for more complex boundary conditions, such as rugged cartilage surfaces sheared against each other. The insights from this work contributes to an understanding at the molecular-level of how lubricin’s conformational dynamics tailor its lubricating function in synovial joints.

## Supporting information

Figures S1--S3 and Tables S1--S10 are included in the Supporting Information.

## Acknowledgement

We gratefully acknowledge support by the German Research Foundation (DFG) through the RTG/GRK-2450 graduate school program; the Klaus Tschira Foundation; the state of Baden-Württemberg through bwHPC. Special thanks are extended to Isabel M Martin, Mohamed Tarek Elewa, and Nicholas Michelarakis for their insightful discussions.

## Supporting Information Available

Figures S1–S3 and Tables S1–S10 are included in the Supporting Information.

## Notes

### Competing Interest Statement

The authors have declared no competing interest.

