## Supplementary material for "O-glycans Expand Lubricin and Attenuate its Viscosity and Shear Thinning": Figures S1--S3 and Tables S1--S10 are included in the Supporting Information.

⊥*Fraunhofer IWM, Wöhlerstraße 11, 79108 Freiburg, Germany*

Table 1: Total number of glycosylated residues in five selected fragments from the central region of lubricin.[2]

| Peptide | Number of glycosylated residues |
|---|---|
| 301-380 | 24 |
| 461-540 | 25 |
| 621-700 | 8 |
| 781-860 | 12 |
| 941-1020 | 23 |

Table 2: Distribution of glycosylated residues and their corresponding five distinct O-glycan types within the 301-380 segment across five different fragments.

| Glycosylation site | O-glycan type | | | | |
|---|---|---|---|---|---|
| | fragment1 | fragment2 | fragment3 | fragment4 | fragment5 |
| 304 | Type4 | Type4 | Type2 | Type4 | Type4 |
| 305 | Type3 | Type3 | Type3 | Type4 | Type3 |
| 306 | Type5 | Type4 | Type4 | Type4 | Type1 |
| 310 | Type1 | Type4 | Type4 | Type3 | Type3 |
| 312 | Type4 | Type4 | Type4 | Type4 | Type6 |
| 316 | Type3 | Type1 | Type3 | Type4 | Type3 |
| 317 | Type1 | Type4 | Type4 | Type3 | Type1 |
| 324 | Type3 | Type4 | Type3 | Type3 | Type3 |
| 325 | Type3 | Type3 | Type3 | Type4 | Type3 |
| 332 | Type4 | Type4 | Type1 | Type6 | Type3 |
| 345 | Type3 | Type3 | Type4 | Type3 | Type4 |
| 346 | Type3 | Type1 | Type3 | Type3 | Type2 |
| 351 | Type3 | Type3 | Type4 | Type4 | Type3 |
| 353 | Type3 | Type2 | Type1 | Type3 | Type4 |
| 354 | Type3 | Type4 | Type3 | Type3 | Type5 |
| 360 | Type3 | Type3 | Type3 | Type4 | Type3 |
| 361 | Type3 | Type4 | Type3 | Type4 | Type4 |
| 362 | Type5 | Type4 | Type4 | Type3 | Type3 |
| 367 | Type3 | Type3 | Type6 | Type3 | Type4 |
| 369 | Type4 | Type1 | Type3 | Type4 | Type4 |
| 370 | Type4 | Type3 | Type4 | Type4 | Type3 |
| 373 | Type3 | Type3 | Type5 | Type3 | Type6 |
| 376 | Type1 | Type3 | Type5 | Type4 | Type3 |
| 377 | Type4 | Type4 | Type4 | Type4 | Type3 |

Table 3: Distribution of glycosylated residues and their corresponding five distinct O-glycan types within the 461-540 segment across five different fragments.

| Glycosylation site | O-glycan type | | | | |
|---|---|---|---|---|---|
| | fragment1 | fragment2 | fragment3 | fragment4 | fragment5 |
| 462 | Type3 | Type4 | Type4 | Type3 | Type3 |
| 463 | Type4 | Type2 | Type4 | Type3 | Type3 |
| 470 | Type1 | Type3 | Type3 | Type1 | Type1 |
| 471 | Type1 | Type3 | Type3 | Type3 | Type4 |
| 477 | Type3 | Type3 | Type4 | Type1 | Type1 |
| 478 | Type2 | Type1 | Type3 | Type3 | Type3 |
| 485 | Type2 | Type3 | Type2 | Type1 | Type3 |
| 493 | Type6 | Type3 | Type3 | Type4 | Type3 |
| 494 | Type3 | Type3 | Type3 | Type4 | Type1 |
| 501 | Type3 | Type4 | Type4 | Type2 | Type1 |
| 502 | Type1 | Type4 | Type3 | Type5 | Type3 |
| 509 | Type4 | Type4 | Type4 | Type3 | Type4 |
| 510 | Type4 | Type3 | Type3 | Type4 | Type3 |
| 511 | Type3 | Type4 | Type6 | Type5 | Type4 |
| 515 | Type4 | Type3 | Type3 | Type3 | Type4 |
| 517 | Type3 | Type1 | Type5 | Type4 | Type6 |
| 518 | Type3 | Type4 | Type3 | Type3 | Type3 |
| 525 | Type1 | Type6 | Type3 | Type4 | Type6 |
| 526 | Type3 | Type3 | Type4 | Type3 | Type3 |
| 527 | Type3 | Type3 | Type3 | Type4 | Type3 |
| 529 | Type3 | Type1 | Type6 | Type3 | Type4 |
| 532 | Type3 | Type6 | Type4 | Type4 | Type4 |
| 533 | Type3 | Type3 | Type3 | Type3 | Type3 |
| 534 | Type4 | Type3 | Type4 | Type1 | Type3 |
| 540 | Type4 | Type4 | Type4 | Type4 | Type3 |

Table 4: Distribution of glycosylated residues and their corresponding five distinct O-glycan types within the 621-700 segment across five different fragments.

| Glycosylation site | O-glycan type | | | | |
|---|---|---|---|---|---|
| | fragment1 | fragment2 | fragment3 | fragment4 | fragment5 |
| 627 | Type1 | Type3 | Type2 | Type1 | Type1 |
| 676 | Type3 | Type3 | Type1 | Type3 | Type3 |
| 683 | Type4 | Type3 | Type3 | Type3 | Type1 |
| 684 | Type4 | Type3 | Type6 | Type4 | Type4 |
| 691 | Type1 | Type3 | Type4 | Type3 | Type4 |
| 692 | Type3 | Type3 | Type5 | Type3 | Type1 |
| 699 | Type3 | Type3 | Type4 | Type1 | Type1 |
| 700 | Type3 | Type3 | Type3 | Type4 | Type4 |

Table 5: Distribution of glycosylated residues and their corresponding five distinct O-glycan types within the 781-860 segment across five different fragments.

| Glycosylation site | O-glycan type | | | | |
|---|---|---|---|---|---|
| | fragment1 | fragment2 | fragment3 | fragment4 | fragment5 |
| 781 | Type5 | Type3 | Type1 | Type3 | Type6 |
| 784 | Type3 | Type3 | Type3 | Type3 | Type5 |
| 785 | Type3 | Type4 | Type3 | Type3 | Type4 |
| 792 | Type3 | Type3 | Type4 | Type4 | Type4 |
| 793 | Type3 | Type4 | Type1 | Type4 | Type3 |
| 805 | Type3 | Type3 | Type1 | Type3 | Type3 |
| 811 | Type6 | Type3 | Type3 | Type1 | Type3 |
| 812 | Type4 | Type4 | Type3 | Type6 | Type1 |
| 813 | Type3 | Type6 | Type3 | Type4 | Type3 |
| 829 | Type3 | Type3 | Type4 | Type3 | Type1 |
| 837 | Type3 | Type3 | Type1 | Type4 | Type4 |
| 838 | Type3 | Type4 | Type4 | Type3 | Type6 |

Table 6: Distribution of glycosylated residues and their corresponding five distinct O-glycan types within the 941-1020 segment across five different fragments.

| Glycosylation site | O-glycan type | | | | |
|---|---|---|---|---|---|
| | fragment1 | fragment2 | fragment3 | fragment4 | fragment5 |
| 941 | Type4 | Type3 | Type1 | Type3 | Type3 |
| 943 | Type3 | Type5 | Type1 | Type3 | Type3 |
| 944 | Type3 | Type4 | Type1 | Type3 | Type3 |
| 945 | Type4 | Type3 | Type4 | Type3 | Type4 |
| 954 | Type3 | Type1 | Type3 | Type3 | Type3 |
| 956 | Type3 | Type4 | Type3 | Type1 | Type3 |
| 957 | Type6 | Type4 | Type3 | Type4 | Type3 |
| 958 | Type3 | Type3 | Type3 | Type3 | Type3 |
| 961 | Type3 | Type3 | Type4 | Type3 | Type2 |
| 962 | Type1 | Type6 | Type3 | Type3 | Type3 |
| 963 | Type3 | Type2 | Type6 | Type4 | Type1 |
| 964 | Type3 | Type4 | Type4 | Type3 | Type3 |
| 965 | Type4 | Type1 | Type3 | Type3 | Type3 |
| 968 | Type3 | Type3 | Type4 | Type1 | Type3 |
| 975 | Type6 | Type4 | Type3 | Type4 | Type4 |
| 978 | Type1 | Type2 | Type5 | Type3 | Type3 |
| 979 | Type4 | Type3 | Type6 | Type3 | Type1 |
| 980 | Type1 | Type3 | Type3 | Type4 | Type4 |
| 986 | Type4 | Type3 | Type3 | Type6 | Type4 |
| 987 | Type3 | Type3 | Type4 | Type1 | Type3 |
| 988 | Type6 | Type3 | Type3 | Type1 | Type1 |
| 1012 | Type4 | Type3 | Type3 | Type4 | Type1 |
| 1014 | Type3 | Type6 | Type4 | Type3 | Type4 |

Table 7: Total charge for glycosylated and non-glycosylated fragments of different peptides.

| Peptide | Non glycosylated | Glycosylated | | | | |
|---|---|---|---|---|---|---|
| | | fragment1 | fragment2 | fragment3 | fragment4 | fragment5 |
| 301-380 | 5 | -22 | -27 | -27 | -33 | -25 |
| 461-540 | 3 | -25 | -28 | -33 | -26 | -26 |
| 621-700 | -3 | -11 | -11 | -13 | -11 | -10 |
| 781-860 | 3 | -11 | -14 | -8 | -13 | -12 |
| 941-1020 | 6 | -23 | -22 | -22 | -19 | -18 |

Table 8: Simulation details for the equilibrium molecular dynamics of single-chain systems.

| Peptide | Glycosylation | Atoms of peptide (no.) | Atoms of system (no.) | Simulation replicas (no.) | Simulation length (ns) |
|---|---|---|---|---|---|
| **301-380** | non-gly. | 1223 | 0.58M | 3 | 200 |
| | fr.1 | 3443 | 0.66M | | |
| | fr.2 | 3527 | 0.57M | | |
| | fr.3 | 3671 | 0.55M | | |
| | fr.4 | 3791 | 0.92M | | |
| | fr.5 | 3599 | 0.72M | | |
| **461-540** | non-gly. | 1182 | 0.43M | 3 | 200 |
| | fr.1 | 3438 | 0.80M | | |
| | fr.2 | 3594 | 0.78M | | |
| | fr.3 | 3822 | 0.76M | | |
| | fr.4 | 3522 | 0.40M | | |
| | fr.5 | 3522 | 0.60M | | |
| **621-700** | non-gly. | 1182 | 0.56M | 3 | 200 |
| | fr.1 | 1854 | 0.53M | | |
| | fr.2 | 1854 | 0.50M | | |
| | fr.3 | 2022 | 0.52M | | |
| | fr.4 | 1854 | 0.61M | | |
| | fr.5 | 1818 | 0.41M | | |
| **781-860** | non-gly. | 1177 | 0.49M | 3 | 200 |
| | fr.1 | 2353 | 0.50M | | |
| | fr.2 | 2413 | 0.36M | | |
| | fr.3 | 2149 | 0.36M | | |
| | fr.4 | 2377 | 0.40M | | |
| | fr.5 | 2437 | 0.67M | | |
| **941-1020** | non-gly. | 1255 | 0.52M | 3 | 200 |
| | fr.1 | 3547 | 0.72M | | |
| | fr.2 | 3511 | 0.53M | | |
| | fr.3 | 3511 | 0.79M | | |
| | fr.4 | 3307 | 0.71M | | |
| | fr.5 | 3223 | 0.59M | | |

Table 9: Simulation details for the equilibrium molecular dynamics of multi-chain systems.

| Glycosylation | Chains | No. of chains | $\rho$ [kg/m$^3$] | Atoms of system system (no.) | Simulation replicas (no.) | Simulation length (ns) |
|---|---|---|---|---|---|---|
| Non | non-gly. of 301-380 | 30 | 66.5 | 0.83M | 4 | 200 |
| Non | random from all non-gly. fragments | 73 | 158.6 | 0.82M | 4 | 200 |
| Non | non-gly. of 301-380 | 101 | 226.6 | 0.81M | 4 | 200 |
| Medium | random from all fragments | 30 | 160.5 | 0.82M | 4 | 200 |
| Highly | fragment4 of 301-380 | 30 | 230.6 | 0.81M | 4 | 200 |

Table 10: Shear-driven non-equilibrium MD simulation parameters

| deforming speeds nm/ns | Shear rates ns$^{-1}$ | Simulation length (ns) |
|---|---|---|
| 0.5 | 0.03 | 600 |
| 1.0 | 0.06 | 400 |
| 2.0 | 0.13 | 200 |
| 4.0 | 0.25 | 100 |
| 8.0 | 0.51 | 100 |
| 10.0 | 0.63 | 50 |
| 15.0 | 0.95 | 50 |

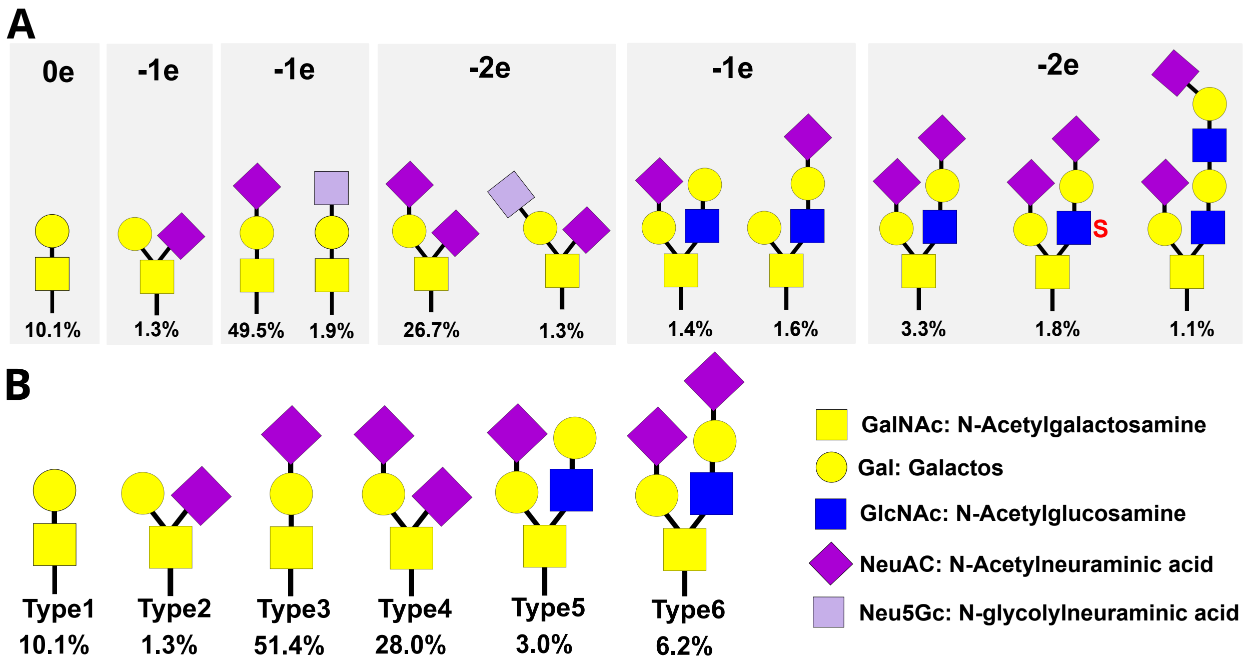


Figure 1: **Composition of O-glycans. A.** All eleven different types of O-glycans of lubricin categorized into six groups based on their structure and charge.[1] **B.** Six selected O-glycans used in our study.

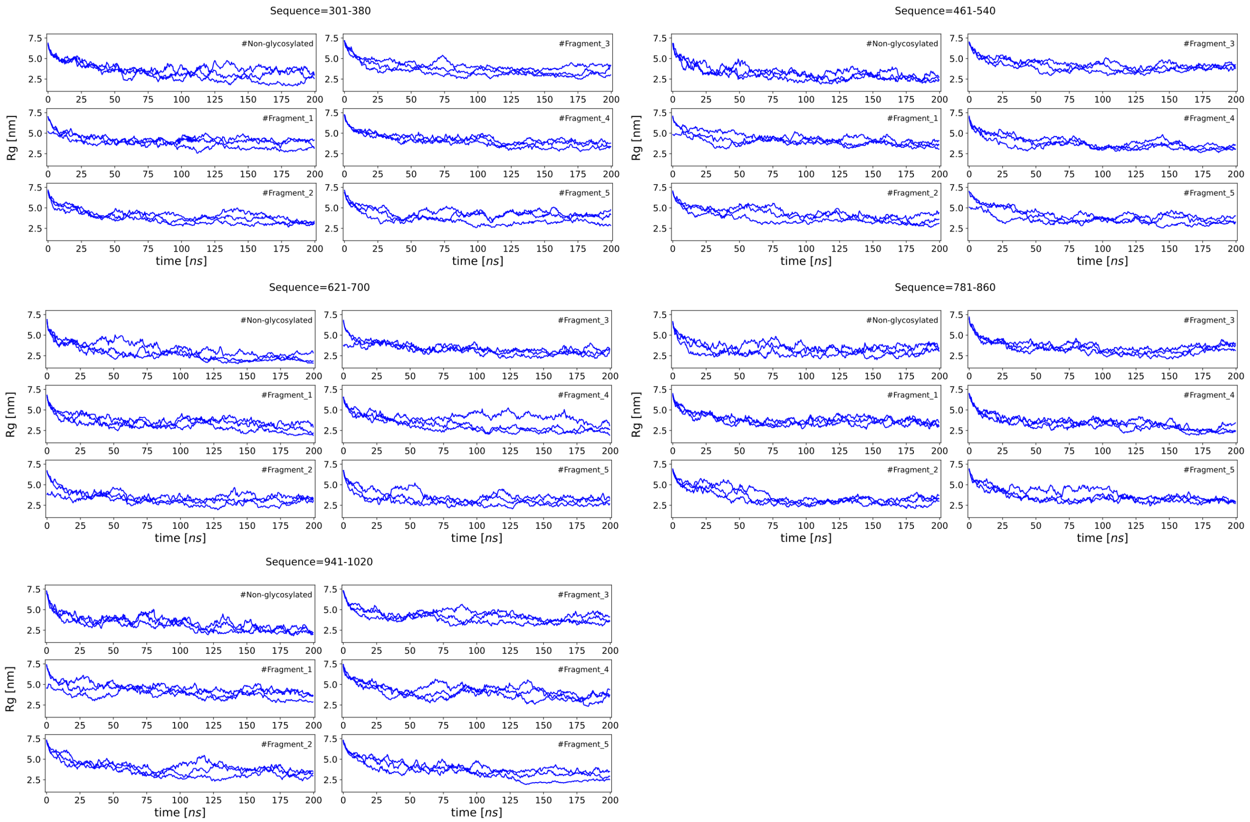


Figure 2: **Radius of gyration time trace for chains with varying levels of glycosylation.** Six different distributions of O-glycans, including one non-glycosylated and five glycosylated chains, were simulated for each of five selected fragments of lubricin. Three replicas for 200 ns were run for each system.

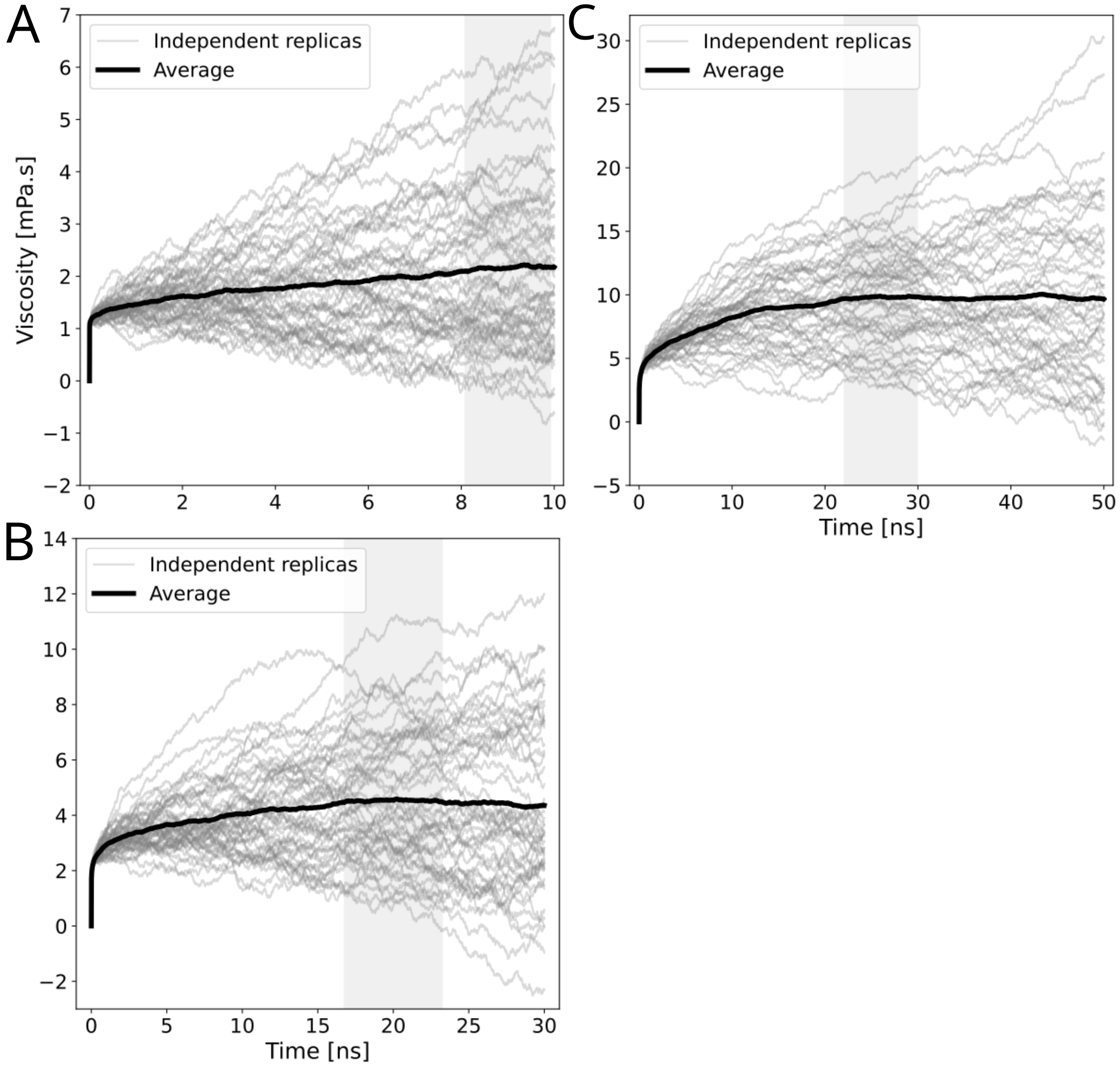


Figure 3: **Zero viscosity comparison in three different systems: A.** 30 non-glycosylated fragments (non-glycosylated system), **B.** 30 random fragments (medium glycosylated system), **C.** 30 glycosylated fragments (highly glycosylated system). Zero-shear viscosity is obtained from the average autocorrelation of all pressure components using the Green-Kubo method. Gray curves represent independent viscosity for each replica (n=50). The black curve displays the average of all these curves. The viscosity was extracted from the plateau-highlighted region, shown in the grey column (average ± standard error).
